# Expression of macromolecular organic nitrogen degrading enzymes identifies potential mediators of soil organic N availability to an annual grass

**DOI:** 10.1101/2020.12.14.422732

**Authors:** Ella T. Sieradzki, Erin E. Nuccio, Jennifer Pett-Ridge, Mary K. Firestone

**Affiliations:** Department of Environmental Science, Policy and Management, University of California, Berkeley, California, USA; Physical and Life Sciences Directorate, Lawrence Livermore National Laboratory, Livermore, California, USA

**Keywords:** Rhizosphere, microbiome, protease, chitinase, functional guilds, metatranscriptome, gene expression, litter

## Abstract

Nitrogen (N) commonly limits terrestrial plant growth partly because most soil-N is present as macromolecular organic compounds and not directly available to plants. Soil microbes degrade these large N-containing substrates to gradually release plant-available inorganic-N throughout the growing season, potentially meeting plant demand. Knowing which microbes are responsible for release of organic N, as well as their spatiotemporal patterns of activity, can enable microbial management strategies that increase plant access to soil-N and reduce dependency on fertilizer-N. We used time-resolved metatranscriptomes to follow taxonomy-resolved differential expression of N-depolymerizing enzymes.

Taxonomic groups show adaptations based on extracellular proteases to specialized habitats in soil characterized by presence (Betaproteobacteria) or absence (Thaumarcheota) of live roots and root detritus (Deltaproteobacteria and Fungi). A similar increase of eukaryotic chitinases near root detritus hints at predation of fungi. Others demonstrate temporal patterns such as increasing expression over time, implying increased competitiveness with substrate depletion (Chloroflexi). Phylotypes from the same genus can have different potential benefits to the plant based on protease expression (e.g., *Janthinobacterium*), which should be considered when designing bioaugmentation. We identify one *Janthinobacterium* phylotype and two Burkholderiales that may be candidates for bioaugmentation near young roots and a *Rhizobacter* which could benefit mature roots.

## Introduction

Plants are commonly limited by nitrogen (N) in temperate soils since their access to the largest soil N pools is constrained by the activity of microbes responsible for ammonification of macromolecular nitrogen and N2 fixation [1–3]. To a large degree, plants depend on microbial degradation of macromolecular forms of N such as proteins and chitin [4–6] and can potentially increase N availability by releasing exudates that stimulate microbial turnover of organic N pools [7–10]. Depolymerization of high molecular weight detrital organic N is a primary rate-limiting step in soil N mineralization [11, 12] and depends on the activity of extracellular enzymes such as lysozyme, protease, chitinase, nuclease and urease. The resulting N monomers are taken up by microorganisms, and roughly 30% of the amino-acid carbon is respired, leading to excretion of excess ammonium which can benefit the plant [13–16]. While the bulk-scale activity of extracellular N degrading enzymes in soil is established, the spatial and temporal dynamics of gene expression underpinning N mineralization are unknown. Additionally, soil microorganisms with the genomic capacity for organic N degradation are diverse [17], and it is unclear whether there is niche partitioning amongst them [18].

Patterns and controls of soil N depolymerization and mineralization may be highly dependent on the habitat (e.g., rhizosphere, detritusphere), since availability of organic-N substrates varies in both time and space, as does the prevalence of fungal, faunal, and bacterial degraders, which can enable or limit N-cycling. In the rhizosphere, organic N is available as amino acids, nucleotides, niacin, and choline (derived from plant exudates [10]), lignoproteins and aromatics (from sloughed off cells), nucleic acids, and microbial cell wall amino sugar polymers (N-acetylglucosamine, N-acetylmuramic acid) derived from the bloom of cells that develops as roots grow [18]. Prior work indicates that root exudates can increase degradation of soil organic matter by up to 380% [19]. In contrast, the detritusphere has a relatively higher proportion of aromatic N and lignoproteins. Bulk protease enzyme activity has been shown to be enhanced by both litter addition [20] and root exudates [21].

Soil N pools and by proxy, N mineralization activity, change with time [22] in part due to the succession of both exudate quality and microbial communities [10, 23]. At the bulk soil level, activity of extracellular proteases is thought to drive soil N cycling [12], but what controls this activity is not always clear [24]. Subtle distinctions caused by time and soil habitat are likely masked because most studies are conducted on whole soils, which contain a mixture of rhizosphere, detritusphere and bulk regions. In grassland soils, the rhizosphere is a particularly critical hotspot for microbial biomass and activity due to rapid recycling of root exudation and debris, and functions as a quasi-digestive system decomposing molecules inaccessible to plants. However, root exudation, which serves as a carbon source for rhizosphere bacteria, declines near older root sections [25]. Therefore, it has been hypothesized that bacteria in a mature rhizosphere environment may be forced to target less labile, higher C:N sources of organic N such as plant litter [26]. Indeed, it has been shown that the mature *Avena* sp. rhizosphere has higher rates of gross nitrogen mineralization and ammonia consumption compared to the young rhizosphere [28].

To determine spatial and temporal patterns and identify which groups of microbes were primarily responsible for decomposition of protein and chitin, we assessed the expression of genes that code for enzymes degrading macromolecular organic N near actively growing and decomposing roots of a common annual grass *Avena fatua* (wild oat grass). We used wild oat as a potential source for putative beneficial microbes to domestic oat, as wild oat in our system does not experience fertilization and therefore needs to acquire N from soil organic matter via its microbiome. We analyzed 48 metatranscriptomes collected over three weeks of active root growth, using a well-developed plant mesocosm approach[23, 26–28] with a fully-characterized annual grassland soil from Northern California. A different subset of these metatranscriptomes was previously used to analyze expression of genes coding for degradation of carbohydrates, in an effort to determine the effect of roots and litter on carbon cycling [28]. The previous study provided important context that allowed us to compare trends in carbon and nitrogen cycling. The primary goal of the analysis presented here was to discover the roles of indigenous rhizosphere microorganisms in macromolecular nitrogen degradation by extracellular enzymes throughout root growth and aging. This identification is the first step towards *in-situ* inoculation. Inoculation of seeds with plant growth promoting bacteria has shown promise in reducing the need for fertilizers[29] and improving plant biomass[30] as well as rhizosphere available nitrogen[31].

## Materials and methods

### Experimental design, sample collection and sequencing

The experimental design, sample collection and sequence data processing are described in detail in [28]. Briefly, common wild oat *Avena fatua* was grown for six weeks in rhizobox microcosms containing soil from the Hopland Research and Extension Center (HREC) in northern California, a Bearwallow–Hellman loam (pH 5.6, 2% total C, include 7^th^ approx. name here) packed at field bulk density. Roots were then allowed to grow into a sidecar soil region with a transparent wall, where the root growth timeline was marked at 3 days, 6 days, 12 days and 22 days. In half of the sidecars, the soil was amended with dried *A. fatua* root detritus (C:N = 13.4) chopped to 1 mm. Bulk soil bags, inaccessible to roots, were placed in each sidecar; a bulk soil treatment amended with root detritus was also included. At each timepoint, three replicate microcosms were destructively harvested for paired rhizosphere and bulk soil, both detritus amended and unamended, yielding a total of 48 samples representing four treatments at four timepoints: rhizosphere, rhizosphere + litter, bulk, and bulk + litter (sup. fig. S1).

Harvested soil (1 g) was placed immediately in Lifeguard Soil Preservation Reagent (MoBio) and processed according to the company protocol. Roots and supernatant were removed, and the soil was stored in −80°C. DNA and RNA were co-extracted using a phenol-chloroform procedure [32, 33] and separated with an AllPrep kit (Qiagen). RNA was DNase treated (TURBO DNase, Thermo-Fisher Scientific), depleted in ribosomal RNA (Ribo-Zero rRNA Removal Kit, Epicentre) and reverse transcribed into cDNA. Metatranscriptomes were sequenced for 48 samples on an Illumina HiSeq 2000 2×150 (TruSeq SBS v3) at the Joint Genome Institute (JGI), in Walnut Creek CA.

### Expression of macromolecular N degrading genes identified in denovo assembled metatranscriptomes

Raw reads were quality-trimmed (Q20) and rRNA and tRNA reads were removed. Library size, evenness, richness and Shannon diversity were comparable between experimental groups with a mean library size of 43 M paired end reads after filtering. In contrast to the single approach used by Nuccio et al. [28], we also *denovo* assembled quality-controlled metatranscriptomic reads into contigs within each sample. Contigs smaller than 200bp were discarded and the remaining contigs from all samples were clustered at 99% identity with cd-hit-est keeping the longest sequence as the cluster representative [34]. Open reading frames (ORFs) were predicted using Prodigal [35]. Extracellular protease ORFs were identified by reciprocal BLAST to extracellular peptidases from the MEROPS database [17, 36]. We acknowledge that this method may generate a conservative estimate of the number of proteases detected. ORFs were assigned a peptidase group (serine-, metallo-, cysteine-peptidase and others) by their best reciprocal BLAST hit. Taxonomy was determined by best BLAST hit to the NCBI non-redundant database (NR, access date July 29^th^ 2019). Chitinase ORFs were identified using six chitinase hidden Markov models (HMMs) from the Kyoto Encyclopedia of Genes and Genomes (KEGG): K01183, K03791, K13381, K17523, K17524 and K17525. Only the first three were detected in our dataset. Lysozyme ORFs were identified using the HMM for KEGG orthology K07273, extracellular nuclease with K01150 (codes for Dns gene, undetected) and K07004, and urease subunit A, B and C with K01428, K01429 and K01430 respectively. Reads were then mapped back to ORFs requiring minimum identity 95% and 75% breadth using bbmap [37]. Read counts were normalized using DESeq2 [38]. Heat maps of normalized counts were generated in R using the heatmap2 function in gplots [39]. Normalized transcript counts per gene per time point were compared between groups (rhizosphere, litter and litter-amended rhizosphere) using ANOVA and Tukey post-hoc test. Boxplots were generated in ggplot2 [40].

### Expression of extracellular protease genes from a curated collection of Hopland-soil genomes

The quality-controlled reads were mapped using BBsplit [37] requiring 80% identity against a dereplicated reference database of 282 HREC soil genomes including isolates [10], single amplified genomes (SAGs) [28], metagenomic assembled genomes (MAGs) (NCBI PRJNA517182) and stable isotope probing MAGs (SIP-MAGs) [41]. On average, 12.3% (range 6.2-31.9%) of the reads per library mapped unambiguously to genomes from the reference database. This additional approach was taken to investigate gene expression within the context of a genome and to search for overlap in guild membership between extracellular protease defined guilds and previously defined CAZy guilds [28]. Three MAGs that were classified as unknown or domain bacteria in previous studies were assigned taxonomy using GTDB-Tk version 1.3.0 [42, 43]. Verification of the taxonomic assignment of MAG Burkholderiales_62_29 was done using GToTree [44] with complete reference genomes of Betaproteobacteria from RefSeq (Feb 28, 2020).

Open reading frames (ORFs) were predicted in all genomes using prodigal [35] and annotated using KEGG [45] and ggKbase (http://ggkbase.berkeley.edu). Extracellular proteases, which do not have hidden Markov models (HMMs) capable of separating them from intracellular proteases, were identified by gene nucleotide identity of at least 90% and coverage of at least 60% to *de novo* assembled ORFs of extracellular protease from the metatranscriptomes. Gene counts were identified using Rsubread featureCounts [46].

Differential expression of chitinases in the genome collection was already performed in a previous publication on expression of carbohydrate active enzymes (CAZy) and was therefore not repeated here [28].

### Statistical analyses

All features and their abundance (represented by metatranscriptomic read counts normalized to sequencing depth) were analyzed with DESeq2 [38] requiring p-value < 0.05 (adjusted for multiple comparisons). Ordination and visualization were conducted in R using ggplot2 [40] and vegan [47]; PERMANOVA (vegan function adonis) was used to detect significant treatment factors affecting expression of nitrogen depolymerization genes (protease expression ∼ location * treatment * time). We define ‘guilds’ as groups of taxa with similar gene upregulation patterns of extracellular proteases in both time and soil habitat. Guilds were assigned by hierarchical clustering based on differential expression of extracellular proteases compared to unamended bulk soil, generating four functional guilds: “Rhizosphere”, “Detritusphere”, “Aging root”, and “Low response”. Effects of the four experimental treatments (bulk, rhizosphere, litter and litter-amended rhizosphere) on protease gene expression were assessed by ANOVA (p adjusted for multiple comparisons), N=3 per time point.

### Data availability

The R code used in this work is publicly available at https://github.com/ellasiera/Protease_ISME_2022. All data used in this publication, including raw reads and genome collection, is publicly available. Metagenomes assembled MAGs can be found in NCBI PRJNA517182, stable isotope probing MAGs in http://ggkbase.berkeley.edu/, single amplified genomes in IMG under study name Mediterranean Grassland Soil Metagenome and single amplified genomes in IMG, see sup. table S2 in Nuccio et al., 2020 for accession numbers [28]. Raw reads can be found in JGI IMG. For JGI ID’s (accession numbers) see supplemental table S1 in Nuccio et al. 2020 [28].

## Results

### Expression of enzymes targeting macromolecular N

Extracellular enzymes depolymerizing macromolecular N in soil mainly target proteins, cell wall components, and nucleic acids[48]. Multiple genes for these functions were identified in this analysis: extracellular nuclease (Xds), urease (UreABC), lysozyme targeting peptidoglycan, chitinase (*chit1*) targeting fungal cell walls and insect exoskeleton and extracellular proteases. However, expression of extracellular proteases was an order of magnitude higher than the other extracellular N degrading enzymes (sup. fig. S2), and like chitinases, was affected by the presence of litter and roots. In contrast, extracellular nuclease (Xds), urease (UreABC) and lysozyme were not influenced by either living or decaying roots, thus, we chose to focus on extracellular proteases and chitinases.

Normalized transcript abundance for 4948 extracellular protease genes from bacteria (4846) and fungi (102) was significantly affected by time (3-way PERMANOVA, F=2.8, p=0.038), litter amendment (F=118.9, p<0.001) as well as interactions between time:treatment (F=26.9, p<0.001), time:location (F=7.7, p<0.001) and location:treatment (F=15.5, p<0.001). Expression in unamended soil (no litter) generally increased slightly over time (fig. 1A), whereas litter-amended soil had initially high expression that then decreased and leveled off over time (fig. 1B).

**Figure 1:**
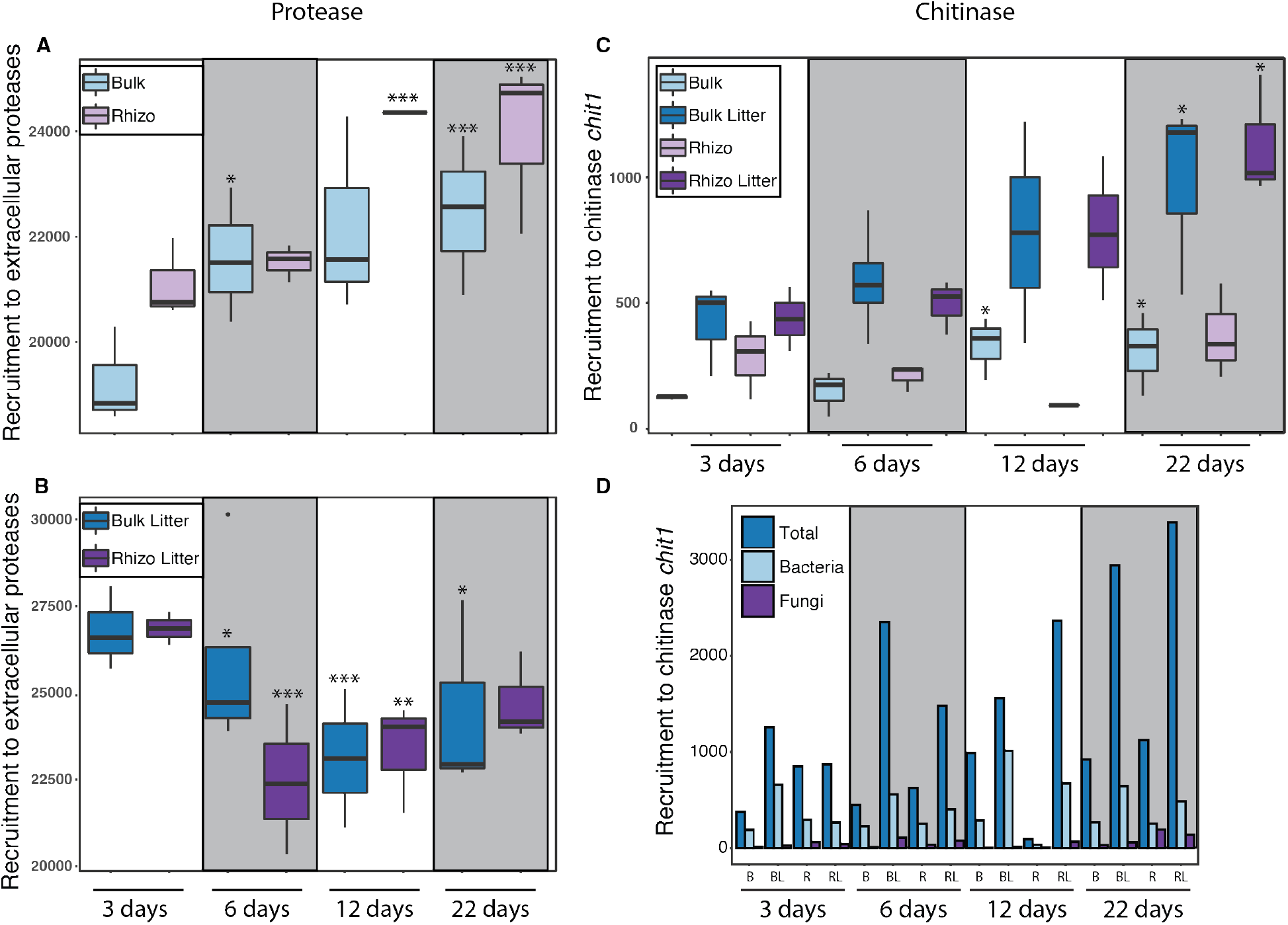
Aggregated expression of extracellular proteases in bulk and rhizosphere soils from common wild oat *Avena fatua* microcosms, grown without root litter amendment (A), or with litter amendment (B); expression of chitinase gene *chit1* (C) and mean counts of chitinases at the domain level (D) for unamended bulk soil (B), litter-amended bulk soil (BL), unamended rhizosphere (R) and litter-amended rhizosphere (R L). Boxplots represent 75% of the data, whiskers denote 90% of the data and dots (e.g., panel A bulk litter 6 days) represent outliers. Note that the scale of the Y axis varies between panels. Statistical significance of linear regression compared to 3 days within the same location and treatment group: * p value <0.05, ** p value <0.01, *** p value < 0.001.

The 73 distinct chitinase transcripts were expressed at a substantially lower level than the extracellular proteases. The expression of the *chit1* gene increased over time in litter-amended soils and was higher than in the unamended treatments (3-way PERMANOVA, p<0.001) (fig. 1C). Expression of fungal chitinases was lower than bacterial chitinases, and both were generally lower than eukaryotic chitinases (fig. 1D). Transcripts for chitinase KEGG orthologs *CHI3L1_2, CHI3L1_4* and *chiA* were not detected at all and the putative chitinase K03791 had low expression and no significant effects with time or treatment (data not shown). Chitinase CHID1 was detected but was most closely related to plants and therefore disregarded. A companion study analyzed differential expression of chitinases within our curated genome collection and while only one MAG was found to upregulate chitinase, it did so in the presence of litter [28].

### Structural groups of extracellular proteases

Structural groups of extracellular proteases can be soil-specific and pH dependent [49]. The main groups of extracellular proteases found in our assembled metatranscriptomes were serine-, metallo- and cysteine-proteases (2679, 1949 and 496, respectively out of 5295 variants clustered at 99% identity). Aggregated expression patterns of all variants in each group reflected the same order (serine>metallo>cysteine). Expression of serine-was consistently higher than metallo-protease across all treatments (ANOVA F=2392, p=0) (fig. 2, sup. table S1). Expression of serine-proteases was also significantly greater in the presence of litter compared to no litter (diff=2337, p<0.001), but root litter amendment did not affect expression of metallo- or cysteine-proteases, and there was no significant effect of time or location on these structural groups (sup. table S1).

**Figure 2:**
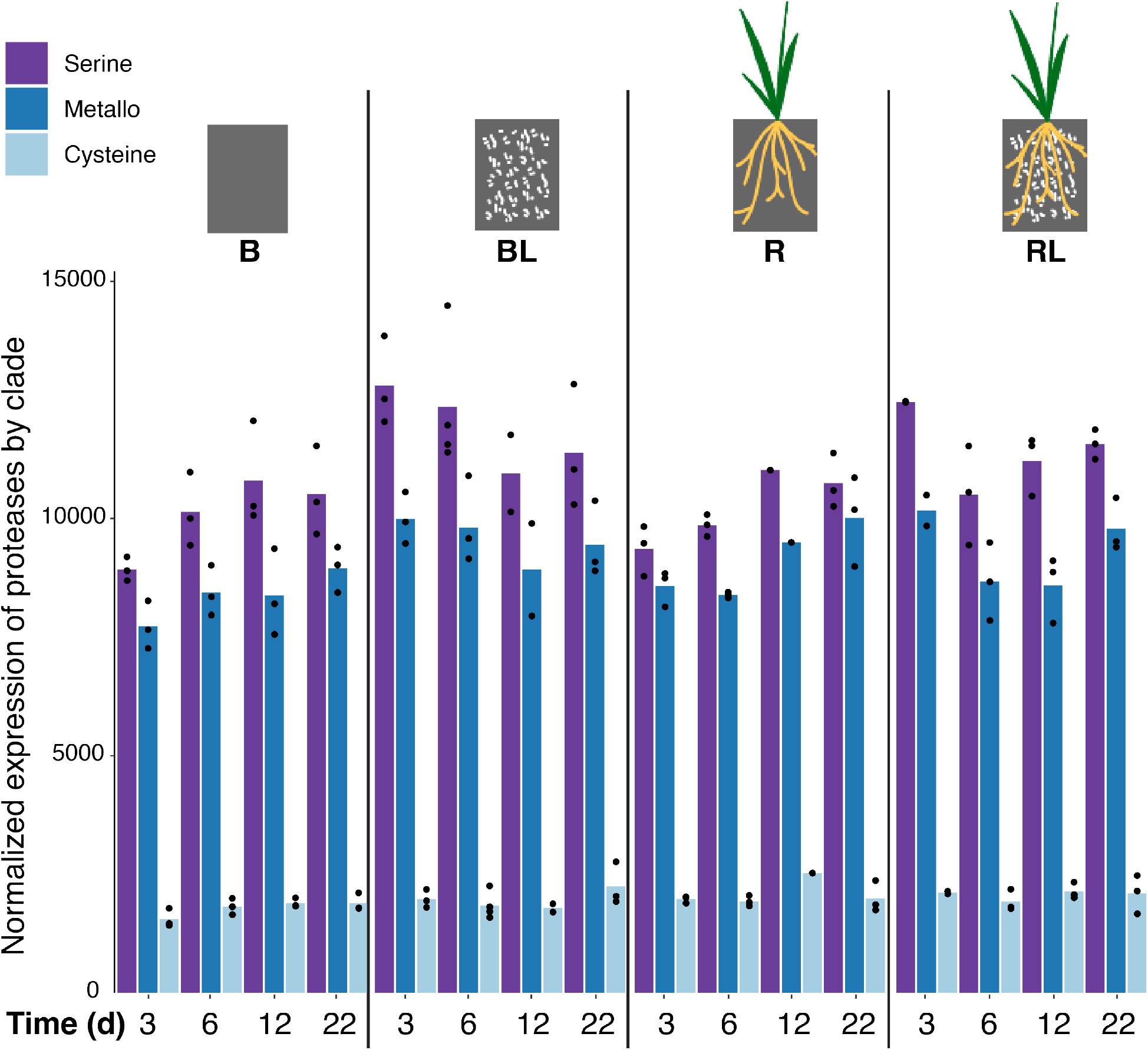
Aggregated normalized expression of extracellular protease structural groups over time in bulk, rhizosphere and root litter-amended soils. Normalized expression per replicate is plotted as dots on top of the bars. Experimental groups shown per time point are (left to right): unamended bulk soil (B), litter-amended bulk soil (BL), unamended rhizosphere (R) and litter-amended rhizosphere (RL). Extracellular protease groups are (left to right): serine-, metallo- and cysteine-protease.

### Taxonomy of extracellular proteases

*De novo* assembled ORFs of extracellular proteases were taxonomically assigned by BLASTP best hit against the NCBI non-redundant database (NR). Since extracellular proteases are not marker genes, we considered taxonomic assignments only at the order level or higher, with the exception of *Rhizobacter*, for which amino acid percent identity values were extremely high (84±8.4% amino acid identity).

In the rhizosphere, Betaproteobacteria proteases were significantly upregulated (ANOVA, p<0.05) (fig. 3A). Of 547 variants of proteases from this class, 442 were assigned as Burkholderiales. In contrast, at most timepoints, proteases of Cyanobacteria and Thaumarchaeota were significantly upregulated in bulk soil compared to the rhizosphere (fig. 3B; sup. table S2). In multiple other taxonomic groups, proteases were significantly upregulated only in the presence of litter: Fungi (fig. 3C), class Deltaproteobacteria (fig. 3D), as well as highly represented phyla Bacterioidetes and Verrucomicrobia (sup. fig. S2; sup. table S2) and classes Chitinophagia and Gammaproteobacteria (sup. table S3). Interestingly, protease expression declined with time for the predatory bacteria Myxococcales (fig. 3E), Bdellovibrionales and Cytophagia, while clades such as Chloroflexi (fig. 3F) and Actinobacteria had increased protease expression at the final sampling point. While the Actinobacteria and Acidobacteria had a high number of variants and high expression of proteases, we did not detect a significant effect of either time or soil habitat/ amendment. Normalized protease expression data are summarized by phylum (sup. fig. S3), class (sup. fig. S4) and order (sup. fig. S5), and ANOVA F and p values are in sup. table S2, sup. table S3 and sup. table S4 respectively. Note that as expression levels varied over taxonomic groups and to emphasize patterns, the y axes do not have the same scale.

**Figure 3:**
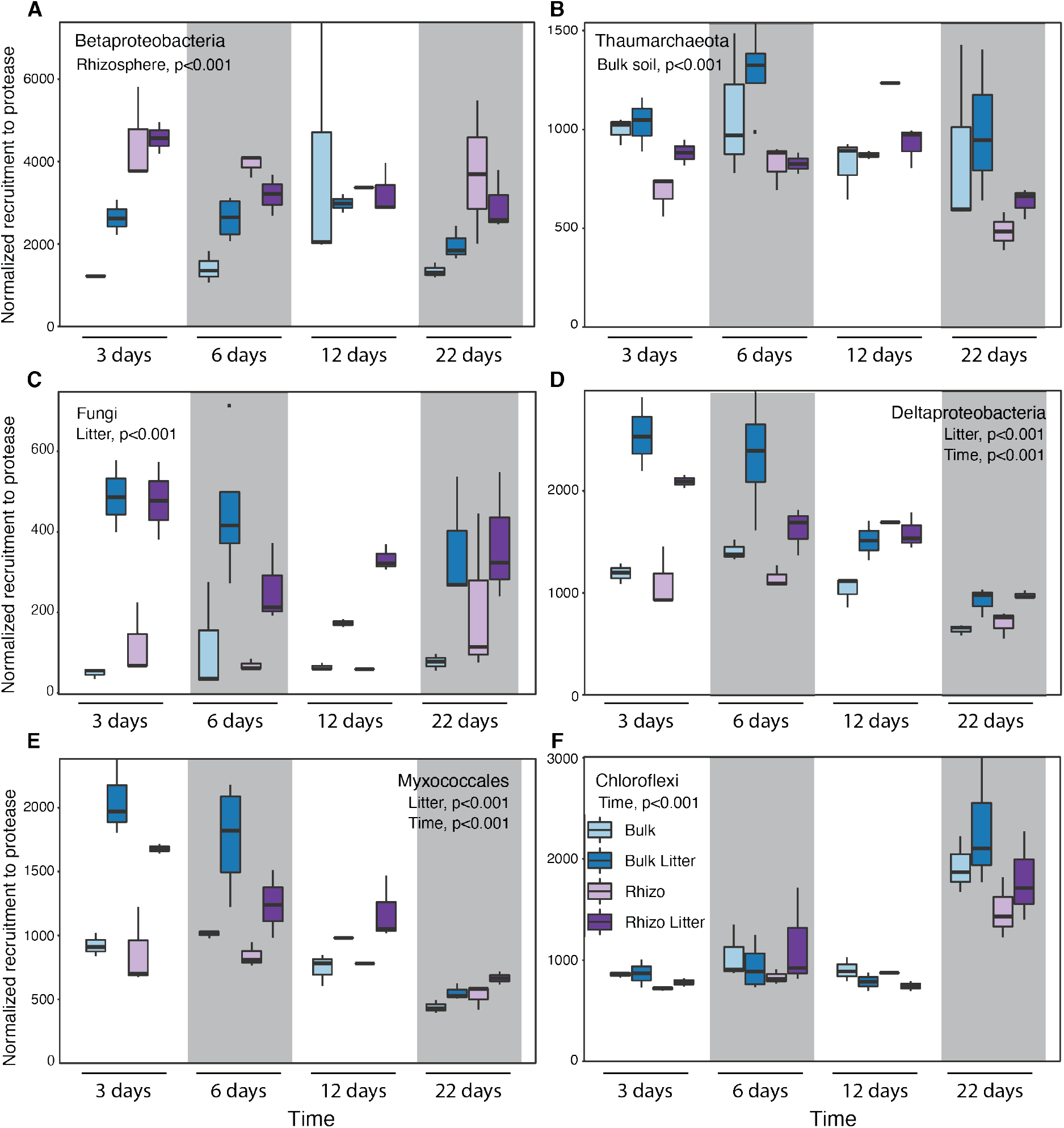
Expression of extracellular proteases over time for select taxonomic groups: (A) Betaprotebacteria (n=547) were upregulated in the rhizosphere, (B) Thaumarchaeota (n=112) were downregulated in the rhizosphere, (C) Fungi (n=99) were upregulated in the presence of litter, (D) Detaproteobacteria (n=228) were upregulated in the presence of litter, (E) Myxococcales (n=147) were upregulated in the presence of litter and decreased over time and (F) Chloroflexi (n=153) increased in the last time point. The legend in panel F applies to all panels.

### Functional guilds

To define functional guilds with a population-centric analysis, we mapped transcriptome reads to a collection of 282 genomes and metagenome-assembled genomes generated from the same soil and site[28]. Read counts were used to determine differential expression compared to unamended bulk soil. Each genome contained multiple genes coding for extracellular proteases. Hierarchical clustering of the mean differential expression of extracellular protease genes within each genome was used to define four functional guilds: “rhizosphere”, “detritusphere”, “aging root” and “low response” (fig. 4). These guilds include bacteria that upregulate expression of extracellular proteases in the presence of live roots (rhizosphere), dead roots (detritusphere), live roots that are beginning to senesce (aging root) or at low but significant levels without a specific pattern (low response)[28]. Similarly, hierarchical clustering based on differential expression of the most highly upregulated *de novo* assembled ORFs compared to unamended bulk soil revealed clear rhizosphere and detritusphere guilds (sup. fig. S6). There was some overlap between guilds identified here and guilds previously identified based on expression of CAZy genes[28], but the degree of overlap varied by guild (sup. fig. S7).

**Figure 4:**
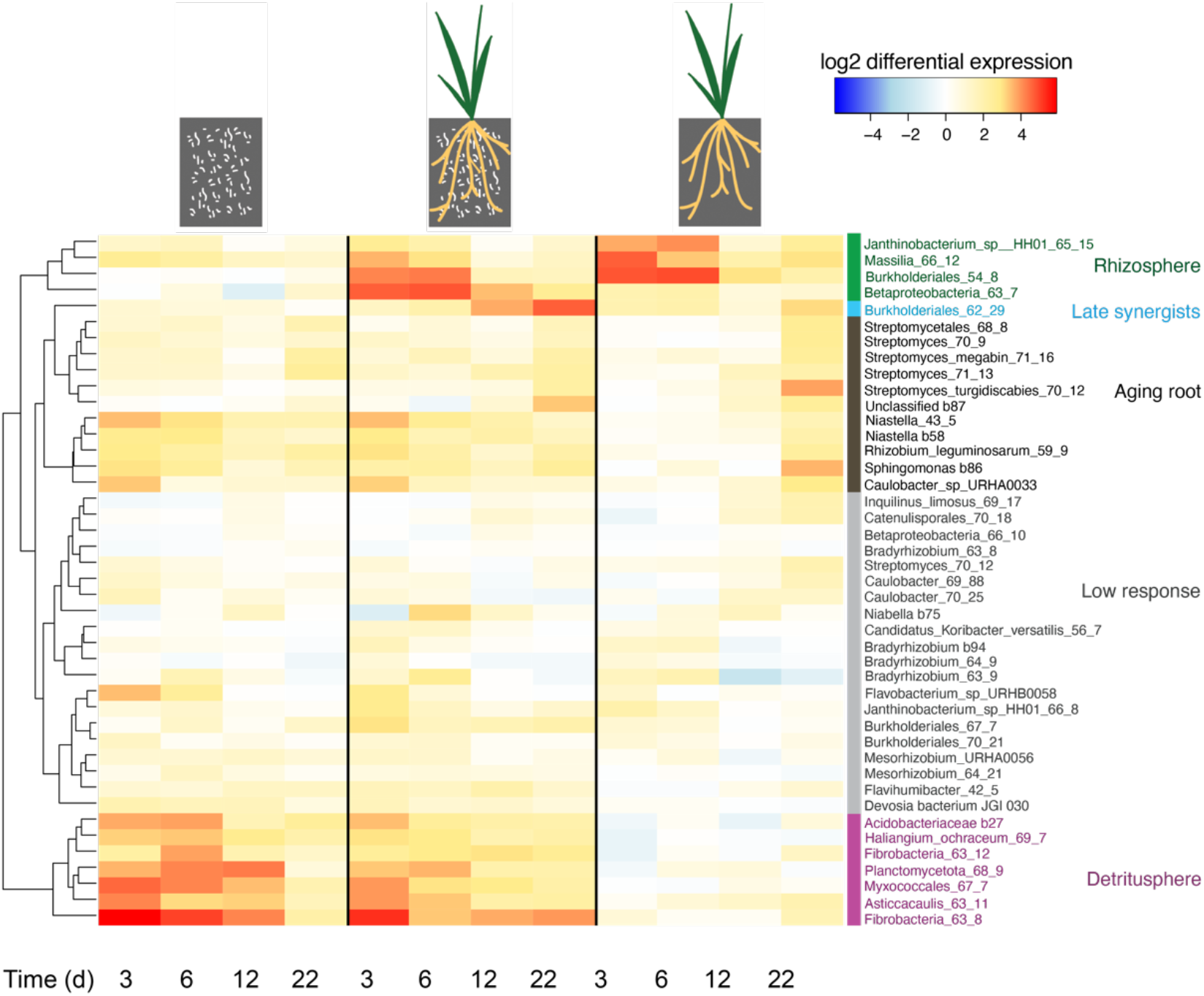
Functional guilds defined by hierarchical clustering of extracellular protease differential gene expression during a 22-day *Avena fatua* microcosm experiment. Each row represents the mean differential expression of extracellular protease genes mapped to a reference genome. Treatments are (left to right): litter-amended bulk soil, litter-amended rhizosphere and unamended rhizosphere. Time points are indicated at the bottom in days. We note that a genome may contain more than one protease gene and that reads were mapped at 80% identity, therefore each genome also represents closely related taxa. Differential expression values per gene that were not statistically significant were converted to zero (0) before averaging.

Within the aging root guild, we noticed one member, representing an aggregated population transcriptome, that increased differential expression of extracellular proteases at the last time point in the presence of both roots and litter more than would have been predicted by the sum of the treatments alone. As this pattern differed from the rest of the guild, we identified this member as a sub-guild labeled “late synergist” due to its expression pattern. This MAG, Burkholderiales_62_29, was identified by two independent phylogenomic analyses as *Rhizobacter*. Burkholderiales_62_29 has 16 different extracellular protease genes, all of which had similar significant upregulation patterns. While the sub-guild here contained only a single *Rhizobacter* MAG, the implication of mapping transcriptomic reads at 80% identity is that each MAG represents a taxonomic “cloud” of at least genus-level diversity. A 16S-rRNA survey of the same samples revealed 10 operational taxonomic units (OTUs) of order Burkholderiales which, like *Rhizobacter*, are not assigned a family [28]. Additionally, this sub-guild may include more taxa for which we have no MAGs and therefore could not be included in this analysis.

## Discussion

Our current understanding of macromolecular nitrogen depolymerization by microbes in soil is based almost entirely on measurement of bulk mineralization rates and genomic surveys. Linking rates and genetic potential is challenging, as rates cannot be broken down by taxonomy and gene presence represents a blueprint which may or may not be acted upon. For example, we show here that of the organic N-degrading enzymes detected in this study, only protease and chitinase are differentially expressed near roots. However, this change in expression was not reflected in changes in the overall microbial community composition [28]. Based on genomic potentials previously reported by Nguyen et al.[17], we hypothesized that expression of structural groups of proteases would be dominated by metalloproteases[17] but expression of serineproteases proved to be significantly higher across experimental groups in this study. Therefore, to identify bacterial candidates that may be beneficial to the plant by improving nitrogen accessibility, analysis of transcripts which provide both taxonomic and metabolic information is crucial.

Chitinase expression, while two orders of magnitude lower than that of proteases, was significantly higher and increased over time in litter-amended samples. This effect could imply that the litter attracted saprotrophic fungi and possibly arthropods, the cell walls of which contain chitin [50]. Indeed, expression of fungal proteases was significantly higher in the presence of litter. As most of the chitinases were most closely related to eukaryotic enzymes, it is likely that the chitin-containing hyphae were preyed upon by soil fauna.

We found that in unamended soil, expression of extracellular proteases was higher in the rhizosphere compared to bulk soil, potentially resulting from input of carbon from root exudates creating a higher demand for nitrogen as well as microbial competition over inorganic nitrogen with the plant. Similarly, DeAngelis et al. showed that protease specific activity was significantly higher in the young rhizosphere compared to bulk soil, but activity (based on enzyme assays) was not significantly different based on root age [27]. In litter-amended soil, expression of extracellular protease was highest regardless of root presence at 3 days, suggesting that litter-added carbon overwhelms the effect of root exudates at this early point in time. At 12 and 22 days there was no difference between litter-amended and unamended soil in either rhizosphere or bulk soil, suggesting that the effect of litter had waned, likely due to substrate degradation. In support, expression of carbohydrate active enzymes after litter addition generally decreased over time [28].

Comparing membership between guilds previously defined by CAZy expression [28] and guilds defined here by hierarchical clustering of extracellular protease expression revealed guild-dependent trends. In the rhizosphere guilds, comprised primarily of Betaproteobacteria known to be commonly associated with rhizosphere soil[23, 51, 52], we found high similarity in membership, whereas in the detritusphere guilds there was low similarity. This implies that in the rhizosphere, the same bacteria break down complex carbon and complex nitrogen, whereas in bulk soil these processes are performed by different taxa. The functional guilds identified here exhibited a pattern of temporal succession throughout plant maturation, where expression of protein degrading enzyme genes in the rhizosphere could be influenced by the host plant through control of exudation, or alternatively, may be a response to competition with the plant for labile nitrogen. Moreover, Nuccio et al. [28], analyzing a different subset of this experimental data, showed that microbial community composition (by 16S-rRNA) in each experimental group changed very little compared to gene expression over the course of this experiment [28]. Therefore, the changing expression patterns over time that we observed are likely related to changes in environmental conditions or cues, such as macromolecular N availability and inorganic N availability, as opposed to wholesale shifts in community composition.

Augmentation of soil or seeds with bacteria that specialize in macromolecular organic N degradation could be used to reduce the use of fertilizers. Ideally, bioaugmentation should use bacteria endemic to the specific plant and soil in order to provide strong competitors[53]. Different patterns of expression of extracellular proteases by various taxonomic groups indicate that potential beneficial partners for the plant change throughout root growth. Hence, functional guild characterization can be useful to decide which taxa should be selected for bioaugmentation, with rhizosphere-, detritusphere- and aging root-specific taxa as candidates. Ideally, organisms from different guilds could be combined into consortia that should effectively contribute more than a single taxon to the nitrogen economy of the plant through mineralization of nitrogen coinciding with plant demand over time.

## Authors’ contributions

ETS performed the data analysis, data interpretation and drafted the manuscript. EEN, JPR and MKF conceived and designed the original rhizosphere-detritusphere study, EEN helped to interpret the data and revise the manuscript. JPR and MKF both interpreted the data and substantially revised the manuscript. All authors approved the submitted version and agreed both to be personally accountable for the author’s own contributions and to ensure that questions related to the accuracy or integrity of any part of the work, even ones in which the author was not personally involved, are appropriately investigated, resolved, and the resolution documented in the literature.

