## Supplemental figures for "Expression of macromolecular organic nitrogen degrading enzymes identifies potential mediators of soil organic N availability to an annual grass"

Supplementary figure S1: Schematic of the experimental design (adapted with permission from Nuccio et al., 2020 26). White boxed represent bulk soil bags made of root-excluding mesh. Dashed black lines aroung the roots denote the rhizosphere.

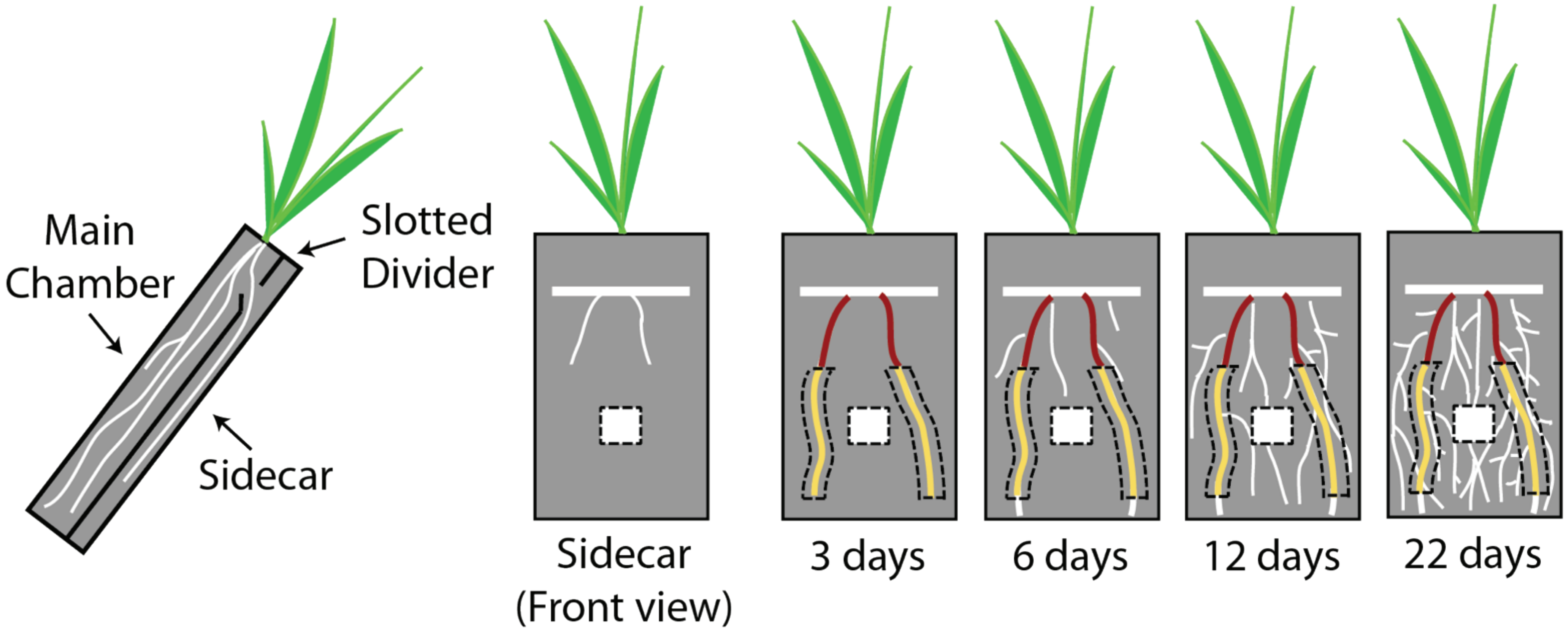

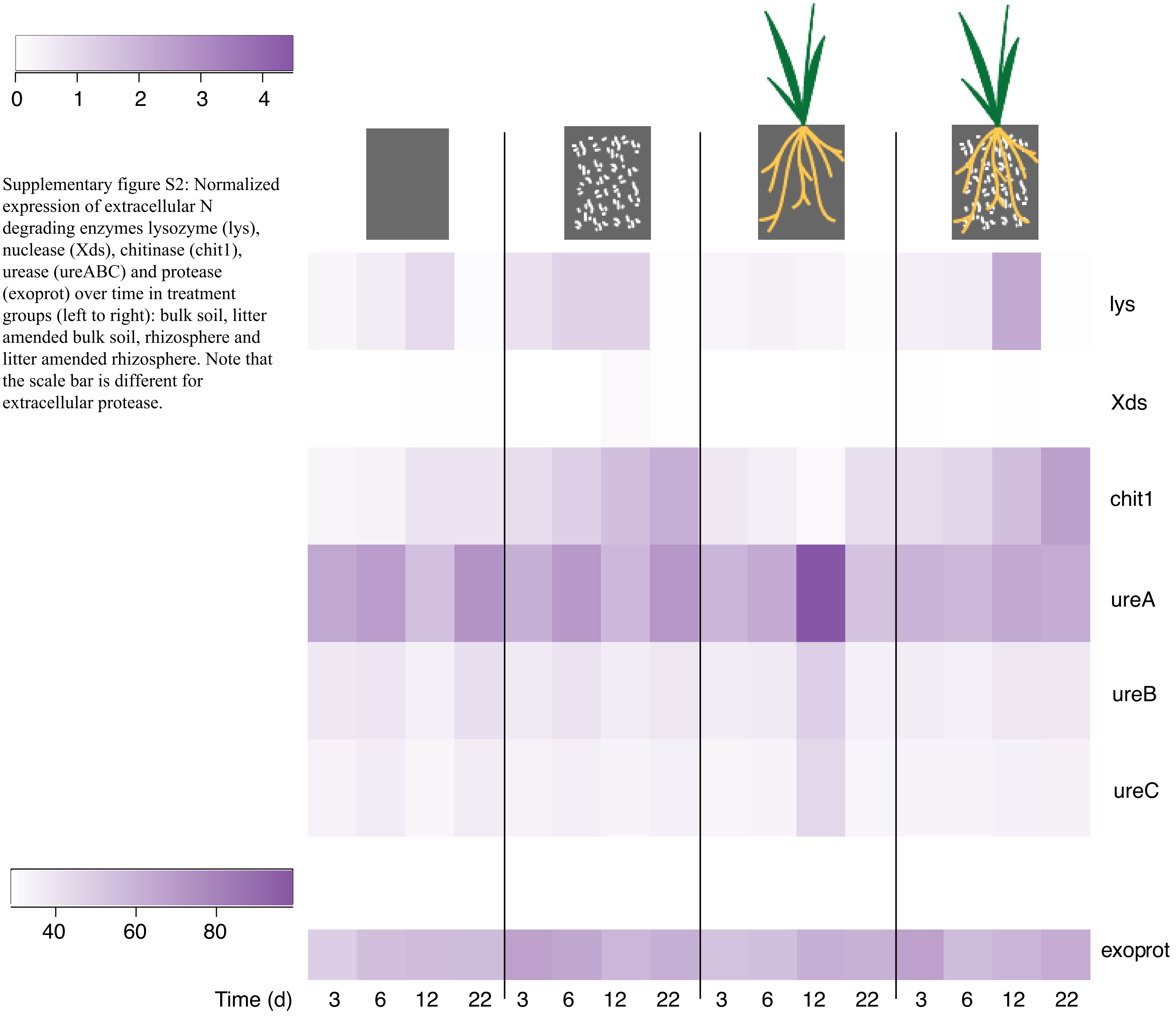


Supplementary figure S2: Normalized expression of extracellular N degrading enzymes lysozyme (lys), nuclease (Xds), chitinase (chit1), urease (ureABC) and protease (exoprot) over time in treatment groups (left to right): bulk soil, litter amended bulk soil, rhizosphere and litter amended rhizosphere. Note that the scale bar is different for extracellular protease.

Supplementary figure S3: Aggregated normalized expression of extracellular proteases by phylum (log transformed) with hierarchical clustering.

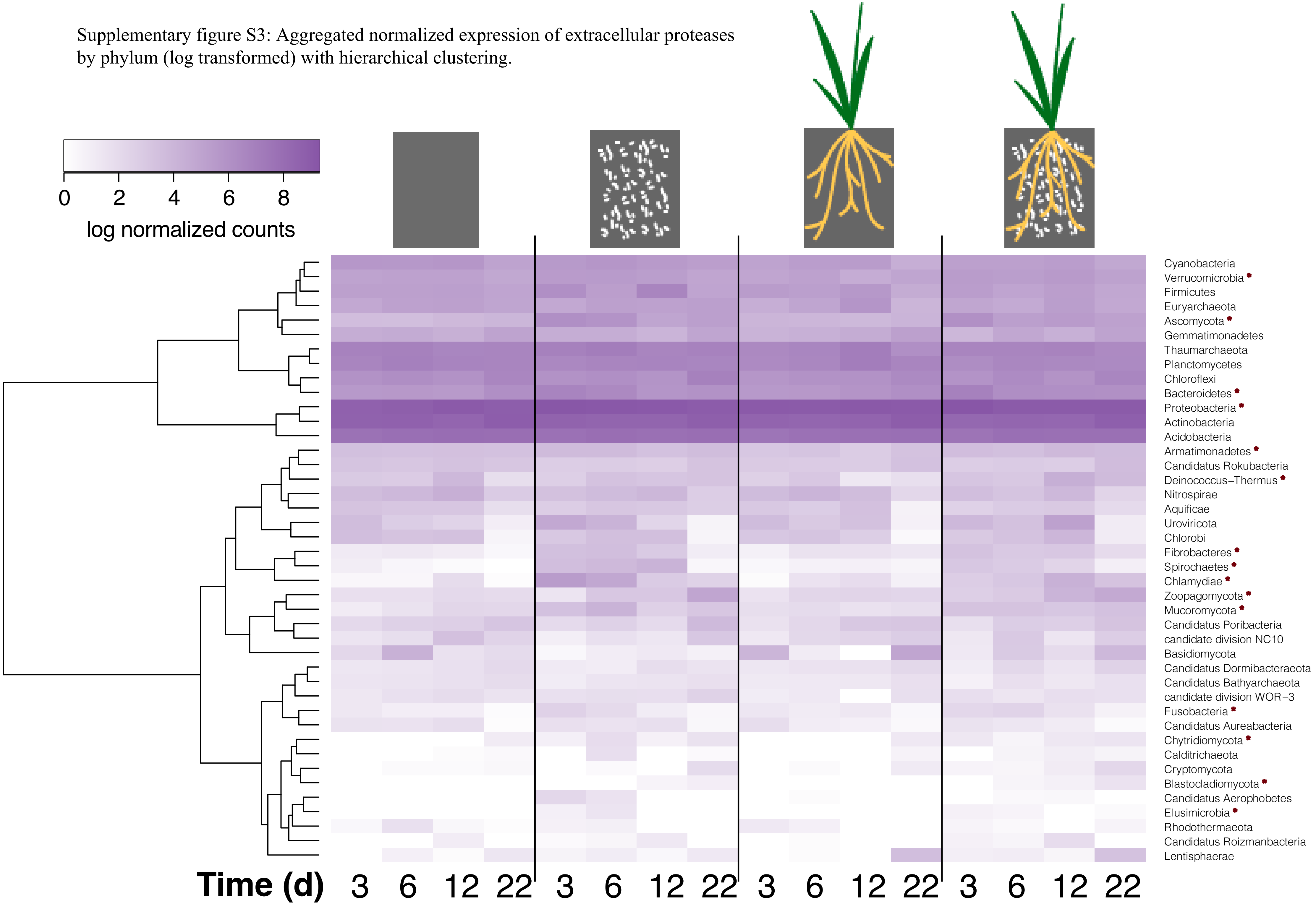

Supplementary figure S4: Aggregated normalized expression of extracellular proteases by class (log transformed) with hierarchical clustering.

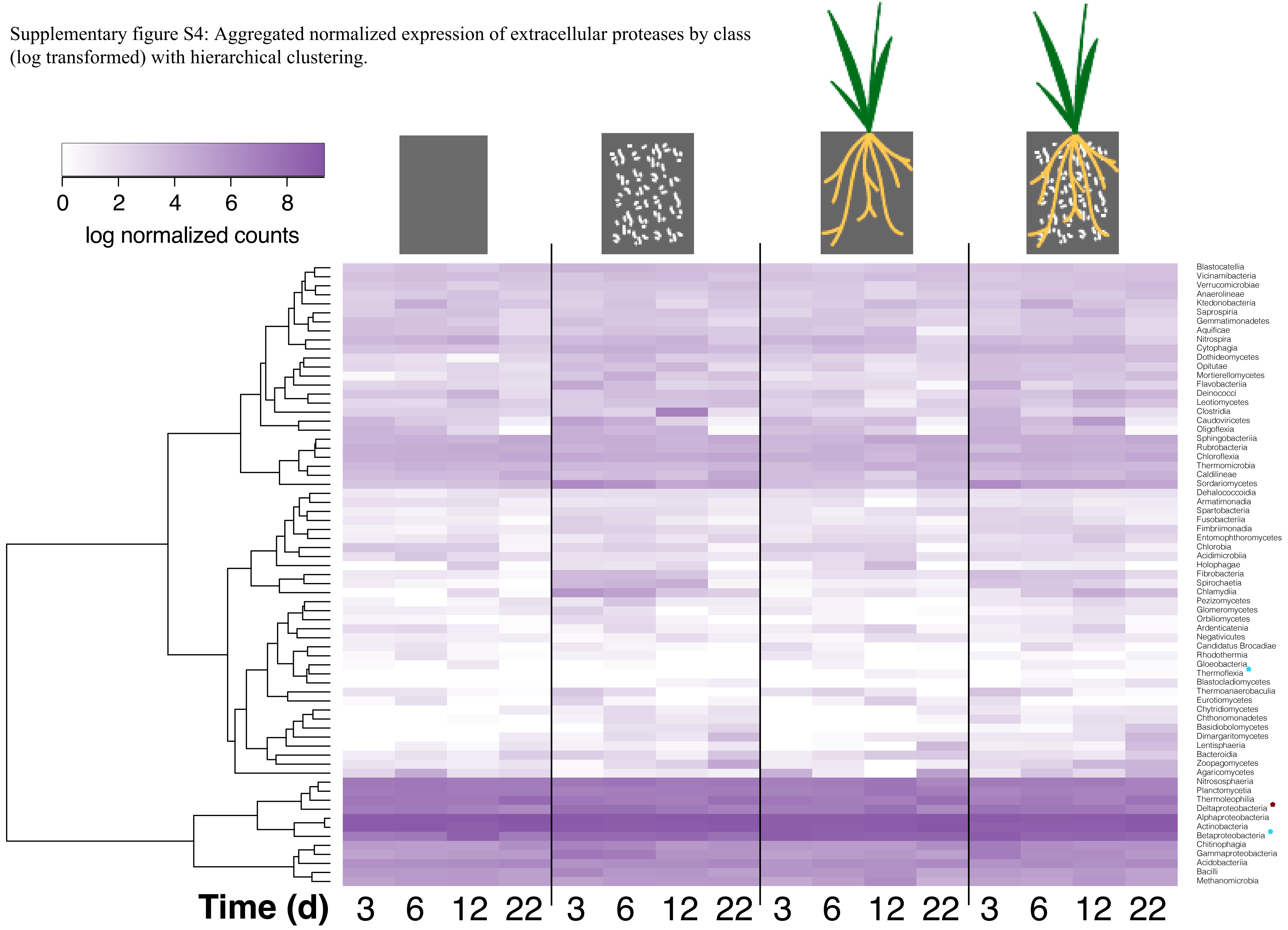

Supplementary figure S5: Aggregated normalized expression of extracellular proteases by order (log transformed) with hierarchical clustering.

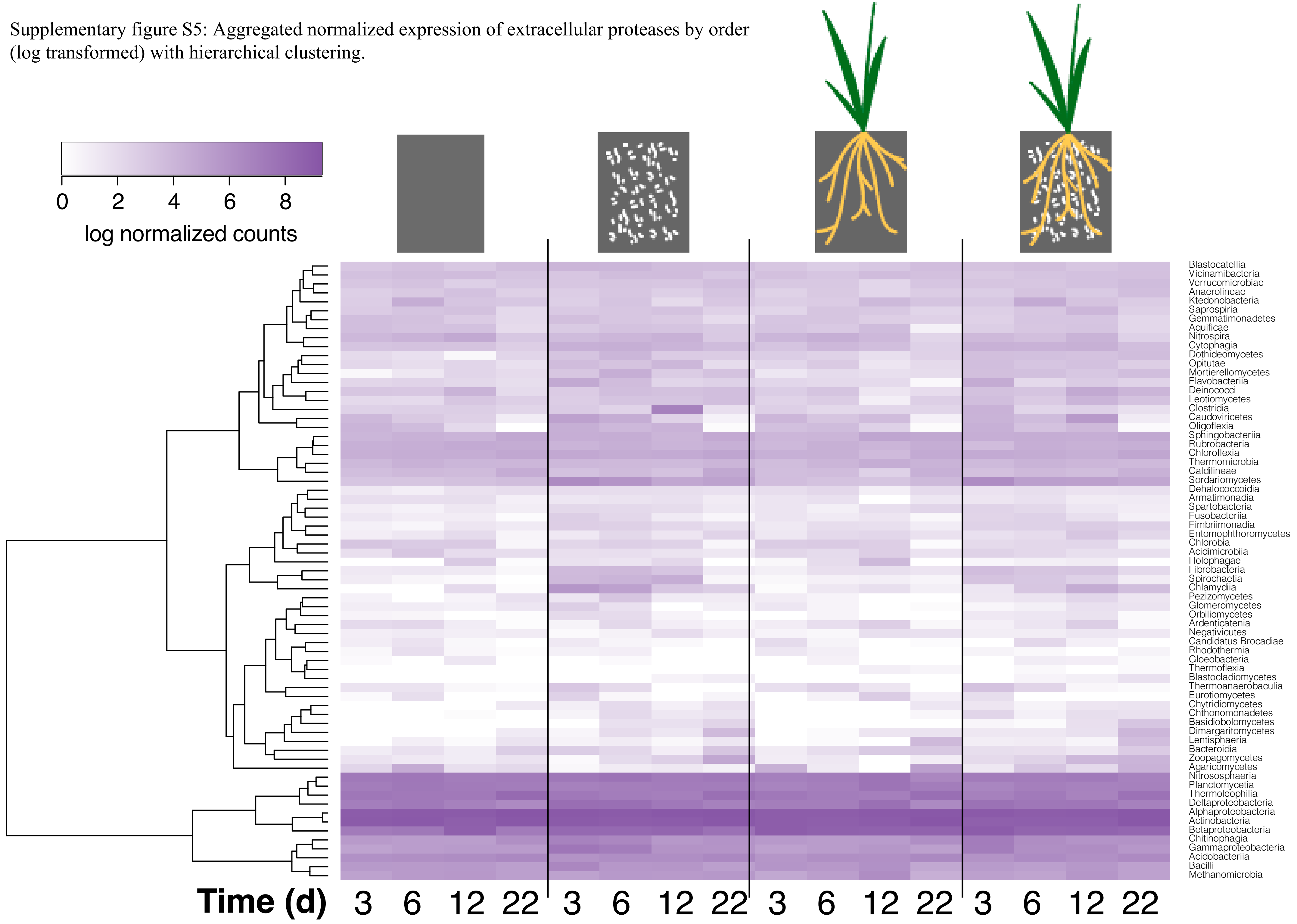

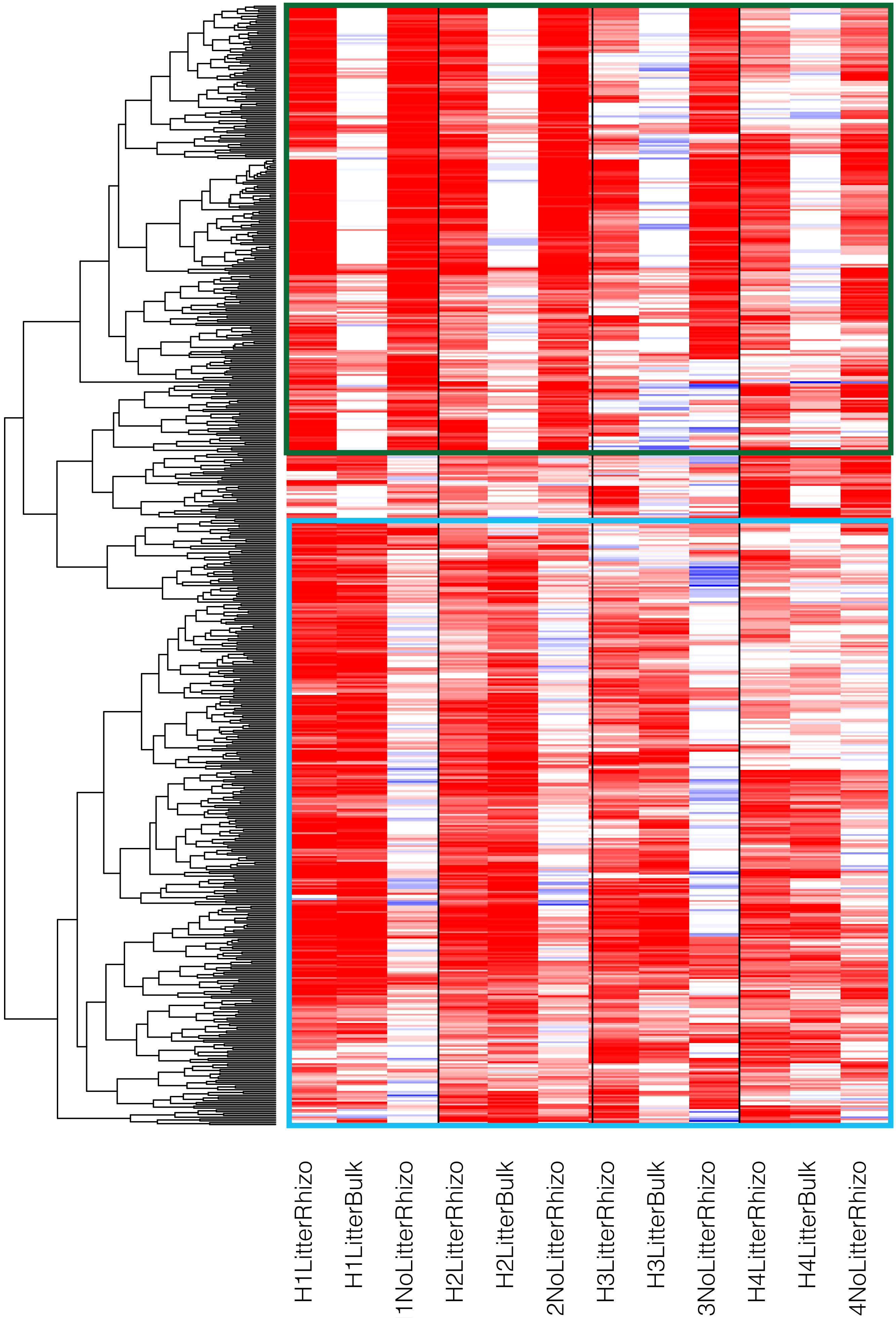


Supplementary figure S6: Functional guilds determined by upregulation of extracellular protease open reading frames assembled from metatranscriptomes. Green = rhizosphere guild, cyan = detritusphere guild

Supplementary figure S7: Number of overlapping genomes between guilds defined by extracellular protease and guilds defined by carbohydrate active enzymes (CAZy).

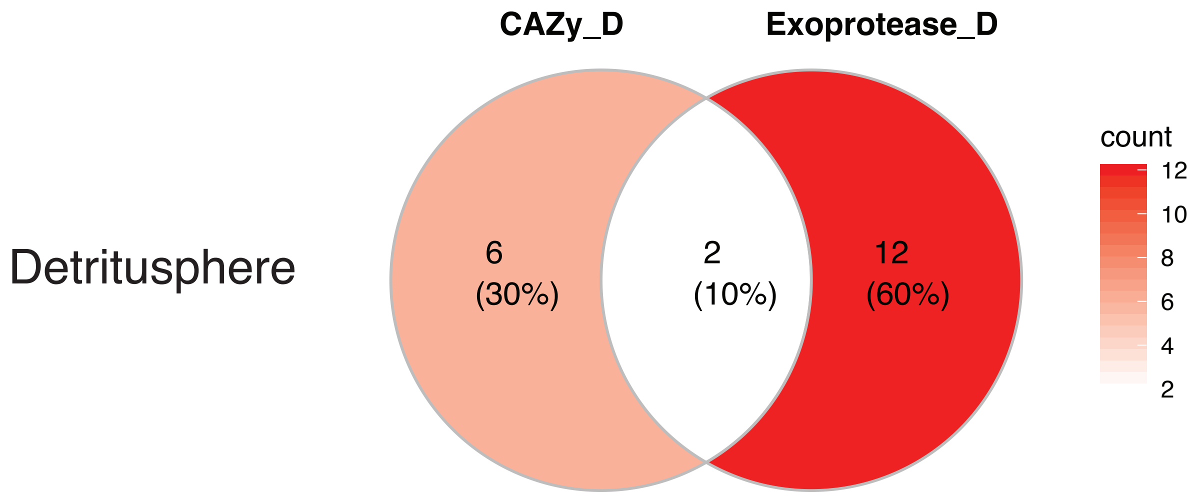


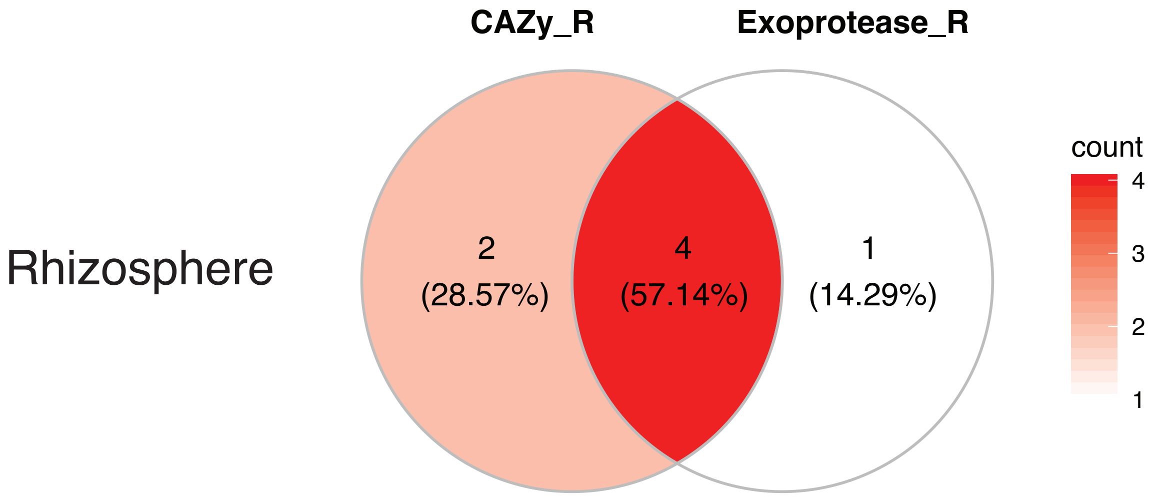


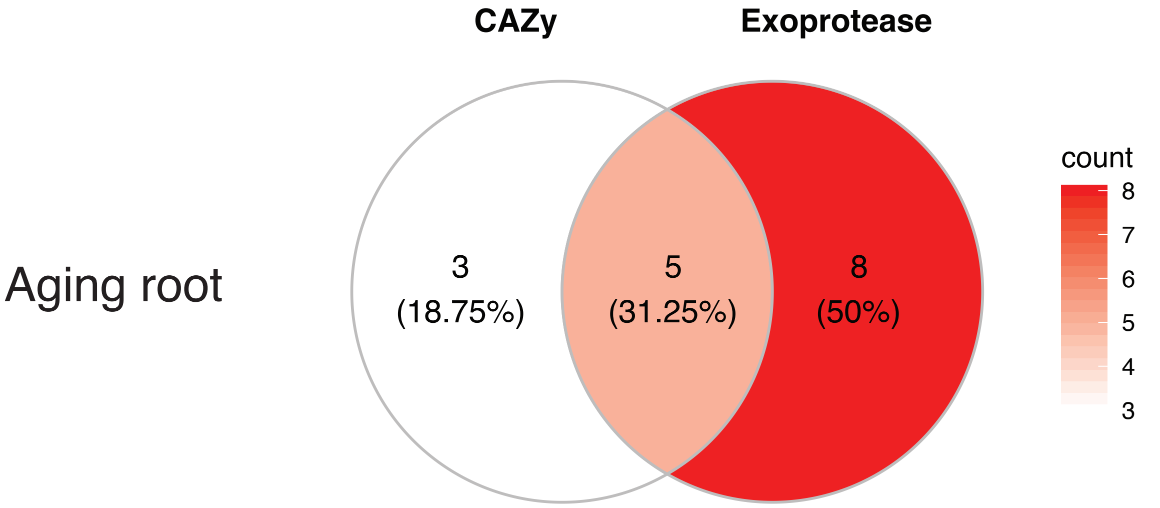
